# *Magnaporthe oryzae* CK2 is involved in rice blast pathogenesis and accumulates in nuclei, nucleoli, at septal and appressoria pores and forms a large ring structure in appressoria

**DOI:** 10.1101/323816

**Authors:** Lianhu Zhang, Dongmei Zhang, Yunyun Chen, Wenyu Ye, Qingyun Lin, Guodong Lu, Daniel J. Ebbole, Stefan Olsson, Zonghua Wang

## Abstract

*Magnaporthe oryzae* (Mo) is a model pathogen causing rice blast resulting in yield and economic losses world-wide. CK2 is a constitutively active, serine/threonine kinase in eukaryotes, having a wide array of known substrates and involved in many cellular processes. We investigated the localization and role of MoCK2 during growth and infection. BLAST search for MoCK2 components and targeted deletion of subunits was combined with protein-GFP fusions to investigate localization. We found one CKa and two CKb subunits of the CK2 holoenzyme. Deletion of the catalytic subunit CKa was not possible and might indicate that such deletions are lethal. The CKb subunits could be deleted but they were both necessary for normal growth and pathogenicity. Localization studies showed that the CK2 holoenzyme needed to be intact for normal localization at septal pores and at appressorium penetration pores. Nuclear localization of CKa was however not dependent on the intact CK2 holoenzyme. In appressoria, CK2 formed a large ring perpendicular to the penetration pore and the ring formation was dependent on the presence of all CK2 subunits. The effects on growth and pathogenicity of deletion of the b subunits combined with the localization indicate that CK2 can have important regulatory functions not only in the nucleus/nucleolus but also at fungal specific structures as septa and appressorial pores.

## INTRODUCTION

Since its discovery (Meggio and Pinna, 2003), the constitutive serine/threonine (S/T) kinase activity of CK2 and the increasing number of proteins it has been shown to phosphorylate have puzzled scientists (Ahmad et al., 2008; Götz and Montenarh, 2016; Meggio and Pinna, 2003). Indeed CK2 has been implicated in a wide range of cellular processes (Götz and Montenarh, 2016). The typical CK2 holoenzyme is a heterotetrametric structure consisting of two catalytic α-units and two regulatory β-subunits (Ahmad et al., 2008). In mammals, there exist two different alpha subunits α (a1) and α’ (a2) and the enzyme can contain any combination of α-subunits (α1α1, α1α2, α2α2) combined with the β-subunits. The CK2 of *Saccharomyces cerevisiae*, also contains two different alpha- and two different β-subunits (b1 and b2) and deletion of both catalytic subunits is lethal (Padmanabha et al., 1990). CK2 functions has been extensively studied in the budding yeast *S. cerevisiae* (Padmanabha et al., 1990), however, functions of CK2 involved in multicellularity might be obscured in yeast. In comparison to yeast, filamentous fungi have many different cell types that allow detailed exploration of cellular differentiation and multicellular development (Shlezinger et al., 2012) and this, in combination with haploid life-cycles, well characterized genomes, and efficient methods for targeted gene replacement, makes fungi like *M. oryzae* As plant pathogens, developmental processes needed for symbiosis can also be explored. We focused our study on *M. oryzae,* one of the most important rice crop pathogens worldwide (Dean et al., 2012).

Our results show that *M. oryzae* CK2 holoenzyme (MoCK2) accumulates in the nucleolus, localizes in structures near septal pores, and assembles to form a large ring structure perpendicular to the appressorium penetration pore and is essential for normal growth and also plant infection.

## MATERIALS AND METHODS

### Material and methods details

#### Fungal strains, culture, and transformation

The *M.oryzae* Ku80 mutant constructed from the wild type Guy11 strain was used as background strain since it lacks non-homologous end joining which facilitates gene targeting (Villalba et al., 2008). Ku80 and its derivative strains (**Table 1**) were all stored on dry sterile filter paper and cultured on complete medium (CM: 0.6% yeast extract, 0.6% casein hydrolysate, 1% sucrose, 1.5% agar) or starch yeast medium (SYM: 0.2% yeast extract, 1% starch, 0.3% sucrose, 1.5% agar) at 25°C. For conidia production, cultures were grown on rice bran medium (1.5% rice bran, 1.5% agar) with constant light at 25°C. Fungal transformants were selected for the appropriate markers inserted by the plasmid vectors. The selective medium contained either 600 μg/ml of hygromycin B or 600 μg/ml of G418 or 50μg/ml chlorimuron ethyl.

**Table 1.**
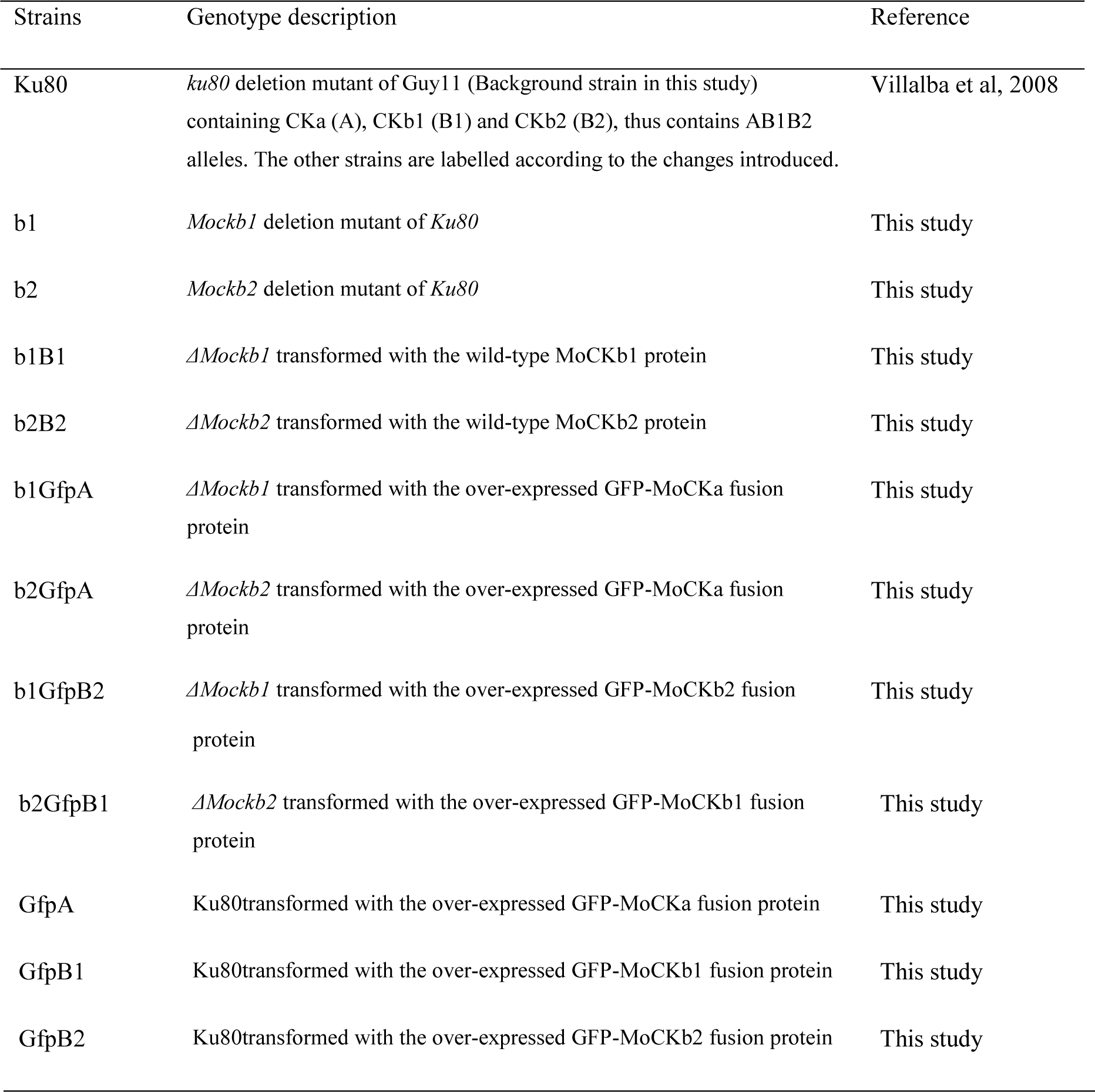
List of strains with simplified strain abbreviations for strains of *M. oryzae* used in this study.

#### MoCKb gene replacement and complementation

Gene replacement mutants of *MoCKb1* encoding protein MoCKb1 were generated by homologous recombination. Briefly, a fragment about 0.9 Kb just upstream of *Mockb1* ORF was amplified with the primers 446AF and 446AR (**Table S1**), so was the 0.7Kb fragment just downstream of *Mockb1* ORF amplified with the primers 446BF and 446BR (**Table S1**). Both fragments were linked with the hygromycin phosphotransferase (*hph*) gene amplified from pCX62 (Zhao et al., 2004) (containing the fragment of TrpC promoter and hygromycin phosphotransferase (*hph*) gene, HPH resistance). Then the fusion fragments were transformed into protoplasts of the background strain Ku80. The positive transformant *ΔMockb1* (strain b1, **Table 1**) was picked from a selective agar medium supplemented with 600 μg/ml of hygromycin B and verified by Southern blot.

For complementation of the mutant, fragments of the native promoter and gene coding region were amplified using the primers 446comF and 446comR listed in **Table S1**. This fragment was inserted into the pCB1532 (Sweigard et al., 1997) to construct the complementation vector using the XbaI and KpnI. Then this vector was transformed into the protoplasts of strain b1. The positive complementation transformant, strain b1B1, was picked up from the selective agar medium supplemented with 50μg/ml chlorimuron ethyl.

As for the *ΔMockb1* deletion mutant, we constructed a knockout vector to delete the *MoCKb2* from the background strain Ku80. All the primers are listed in the **Table S1**. The 1.0Kb fragment upstream of *Mockb2* ORF was amplified with the primers 5651AF and 5651AR, inserted into the plasmid pCX62 using the KpnI and EcoRI to get the pCX-5A vector. The 1.0Kb fragment downstream of *Mockb2* ORF was amplified with the primers 5651BR and 5651BR, inserted into the vector pCX-5A using BamHI and XbaI to construct the knockout vector pCX-5D. Then this vector was transformed into the protoplasts of Ku80. The positive transformants were picked up from the selective medium supplemented with the 600 μg/ml hygromycin B. For complementation of the resulting mutant, strain b2 (**Table 1**), fragments of the native promoter and gene coding region were amplified using the primers 5651comF and 5651comR listed in **Table S1**. This fragment was inserted into pCB1532 to construct the complementation vector using the XbaI and XmaI. Then this vector was transformed into protoplasts of the strain b2. The positive complementation transformant, strain b2B2, was picked up from the selective agar medium supplemented with 50μg/ml chlorimuron ethyl.

#### The construction of localization vectors

In order to detect the localization of MoCK2, we constructed localization vectors. The vector pCB-3696OE containing the RP27 strong promoter (Bourett et al., 2002; Zheng et al., 2007) was used to detect the localization of GFP-MoCKa (strain GfpA). The vector pCB-446OE expressed under RP27 strong promoter was used to detect the localization of GFP-MoCKb1 (strain GfpB1). The vector pCB-5651OE expressed by RP27 strong promoter was used to detect the localization of GFP-MoCKb2 (strain GfpB2).

#### Analysis of conidial morphology, conidial germination and appressoria formation

Conidia were prepared from cultures grown on 4% rice bran medium. Rice bran medium was prepared by boiling 40g rice bran (can be bought for example through Alibaba.com) in 1L DD-water for 30 minutes. After cooling pH was adjusted from to 6.5 using NaOH and 20 g agar (MDL No MFCD00081288) was added before sterilization by autoclaving (121°C for 20 minutes). Conidia morphology was observed using confocal microscopy (Nikon A1^+^). For conidial germination and appressoria formation conidia were incubated on hydrophobic microscope cover glasses (Beckerman and Ebbole, 1996) (Fisherbrand) at 25 °C in the dark. Conidial germination and appressoria formation were examined after 24 h incubation (Beckerman and Ebbole, 1996; Ding et al., 2010).

#### Pathogenicity assay

Plant infection assays were performed on rice leaves. The rice cultivar used for infection assays was CO39. In short, mycelial plugs were put on detached intact leaves or leaves wounded by syringe stabbing. These leaves were incubated in the dark for 24h and transferred into constant light and incubated for 5 days to assess pathogenicity (Talbot et al., 1996). For infections using conidial suspensions (1× 10^5^ conidia/ml in sterile water with 0.02% Tween 20) were sprayed on the rice leaves of 2-week-old seedlings.

### RNA extraction and real-time PCR analysis

RNA was extracted with the RNAiso Plus kit (TaKaRa). First strand cDNA was synthesized with the PrimeScript RT reagent Kit with gDNA Eraser (TaKaRa). For quantitative real-time PCR, *MoCKa, MoCKb1*, and *MoCKb2* were amplified with the primers listed in **Table S1**. β-tubulin (XP_368640) was amplified as an endogenous control. Real-time PCR was performed with the TaKaRa SYBR Premix Ex Taq (Perfect Real Time) (Takara). The relative expression levels were calculated using the 2^-ΔΔCt^ method (Livak and Schmittgen, 2001).

## DATA AND SOFTWARE AVAILABILITY

Alignments, phylogenetic trees, analysis to find orthologues and homologues to well described proteins in other fungal species is available from Figshare either as specific links from the **Table 2** or through this link (LINK TO BE INSERTED) to the Figshare collection specific for this paper (NOTE: a public link to this collection will be inserted here after this paper is accepted and all information in the files will then been made public on Figshare)

**Table 2.** Key resources.

| REAGENT or RESOURCE | SOURCE | IDENTIFIER |
| --- | --- | --- |
| Chemicals, Peptides, and Recombinant Proteins |  |  |
| Hygromycin B | Sangon Biotech | MFCD06795479 |
| G418 | Real-Times (Beijing) Biotechnology Co.,Ltd. | M345810 |
| Chlorimuron ethyl | 9dingchem | 90982-32-4 |
| Agar | MDL | MFCD00081288 |
| Critical Commercial Assays |  |  |
| PCR: RNAiso Plus (TaKaRa) | Takara | 9108 |
| RT-qPCR: TaKaRa SYBR Premix Ex Taq | Takara | RR071A |
| Deposited Data |  |  |
| Phylogenetic trees of *M. oryzae* CK2 components | Figshare, this paper | https://figshare.com/s/751c3d40cb5fda392c20 |
| CKa alignments | Figshare, this paper | https://figshare.com/s/44a9314f00d8dc7c756e |
| CKb1 alignments | Figshare, this paper | https://figshare.com/s/66710db8b62f55d749ee |
| CKb2 alignments | Figshare, this paper | https://figshare.com/s/20104321422b39cad7d5 |
| Experimental Models: Organisms/Strains |  |  |
| *Magnaporthe oryzae*: Ku80 | (Villalba et al., 2008) | N/A |
| Rice, *Oryza sativa* L. *ssp indica* cultivar CO39 | N/A | N/A |
| Strains for CK2 studies in *M. oryzae*, see Table 1 | This paper | N/A |
| Oligonucleotides |  |  |
| Primers for CK2 studies in *M. oryzae*, see Table S1 | This paper | N/A |
| Recombinant DNA |  |  |
| pCX62 | (Zhao et al., 2004) | N/A |
| pCB1532 | (Sweigard et al., 1997) | N/A |
| pCX-5A | This paper | N/A |
| pCX-5D | This paper | N/A |
| pCB-446OE | This paper | N/A |
| pCB-5651OE | This paper | N/A |
| Other |  |  |
| Confocal Microscope | Nikon | A1+ |

### Contacts for reagents and resource sharing

Further information and requests for resources and reagents should be directed to and will be fulfilled by Prof. Stefan Olsson or Prof. Zonghua Wang.

## RESULTS

### Deletion of MoCK2 components

We identified one CKa catalytic subunit ortholog (MoCKa1, MGG_03696) and two MoCKb regulatory subunit orthologs (MoCKb1, MGG_00446 and MoCKb2, MGG_05651) based on BLASTp (NCBI) analysis using protein sequences for the CK2 subunits of *S. cerevisiae,* CKa1, CKa2, CKb1 and CKb2 (Padmanabha et al., 1990) (**Fig. 1A**). Filamentous fungi have just one highly conserved catalytic subunit. In the case of *F. graminearum*, two genes with homology with CKa were identified previously (Wang et al., 2011), one that is highly conserved (we name FgCKa1), and one that is CKa-like (named FgCKa2). It remains to be determined if FgCKa2 is actually a CK2 subunit (See Table 2 for links to Alignments and Phylogenetic trees)

**Fig. 1.**
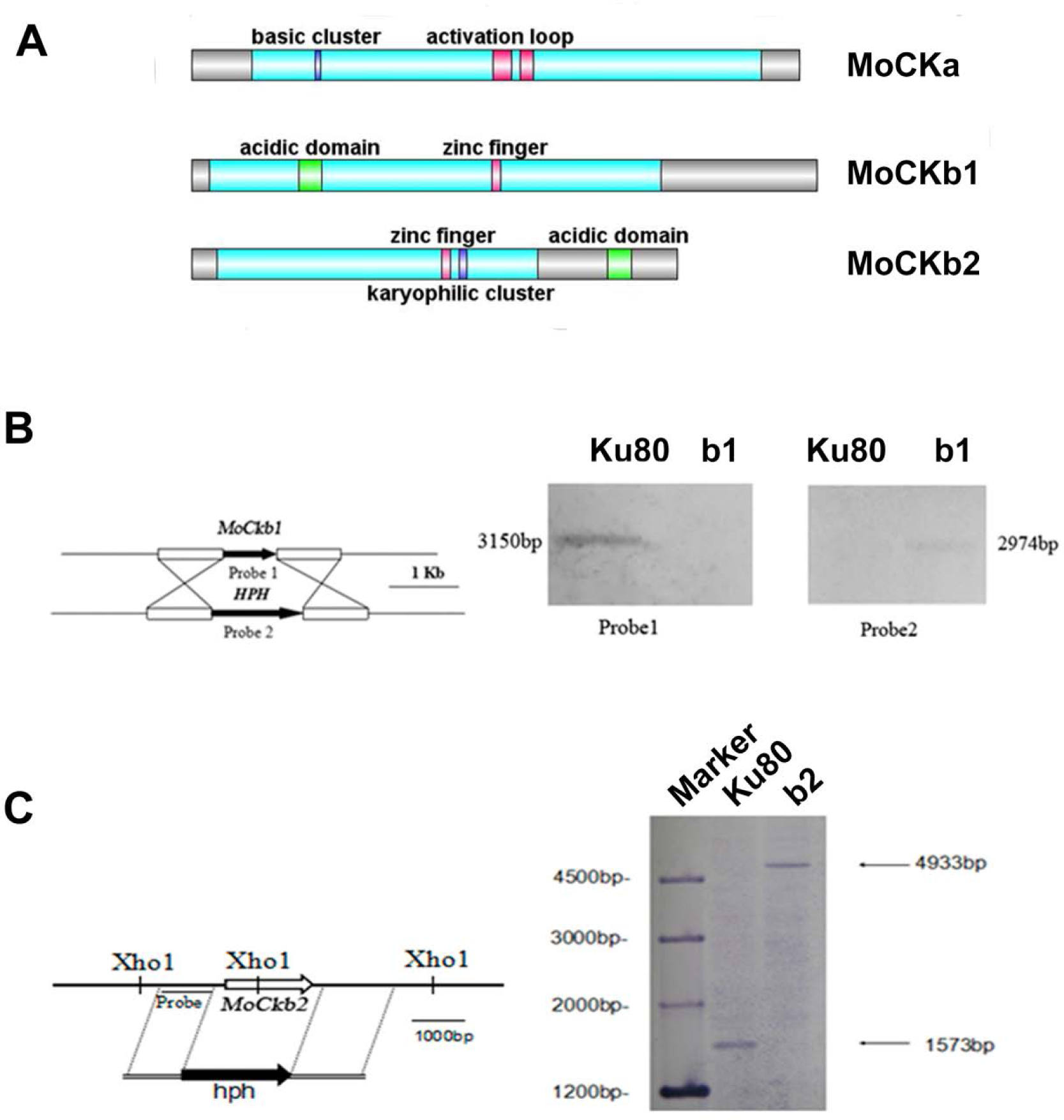
Domain structure of the identified CK2 proteins. **(A).** MoCKa sequence (341 aa) was obtained from NCBI and the 35-320 region contains the protein kinase domain (https://www.ncbi.nlm.nih.gov/nuccore/XM_003716137.1) is labeled light blue, 70-73 is the basic cluster labeled dark blue and functions as a nuclear localization signal (NLS), 170-180 and 185-192 are activation loops (A-loop) labeled red. MoCKb1 sequence (351 aa) was obtained from NCBI and the 11-264 region contains the Casein kinase II regulatory domain (https://www.ncbi.nlm.nih.gov/protein/XP_003718622.1) labeled light blue, 61-74 is the acidic domain labeled green, 169-174 is the zinc finger domain labeled pink. MoCKb2 sequence (273 aa) was obtained from NCBI and the 15-195 region is the Casein kinase II regulatory subunit domain (https://www.ncbi.nlm.nih.gov/protein/XP_003710544.1) labeled light blue, 141-146 is a zinc finger labeled pink, 151-155 is a karyophilic cluster labeled dark blue that functions as a NLS, 234-247 is an acidic domain labeled green. The illustration was made using the DOG 2.0 Visualization of Protein Domain Structures http://dog.biocuckoo.org/. (**B**) *ΔMockb1* mutant (named b1) was verified by Southern blot analysis. The genomic DNA extracted from the strains Ku80 and the mutant *ΔMockb1* was digested with *Nde1* and tested by Southern blot. The different probes ORF and hph (hygromycin fragment) were amplified from the genomic DNA of the background strains Ku80 and the plasmid pCX62 respectively. (**C**) *ΔMockb2* mutant (named b2) was verified by Southern blot analysis. The genomic DNA extracted from the strains *Ku80* and the mutant *ΔMockb2* was digested with *Xho1* and tested by Southern blot. The probe was amplified from the genomic DNA of the background strains Ku80.

Targeted deletions of the two regulatory subunits succeeded (**Fig. S1B, C Fig. 1B,C**). For abbreviations of all strains used in this study see **Table 1**. Attempts to delete the catalytic subunit, CKa, were unsuccessful, consistent with the essential role of CK2 activity (Hermosilla et al., 2005).

The most obvious visible phenotype of the CKb mutants (strain b1 and b2) in culture was reduced growth rate (**Fig. 2A, B**) and conidial morphology in that they produced few conidia and those that were produced had fewer conidial compartments (**Fig. 2C**).

**Fig. 2.**
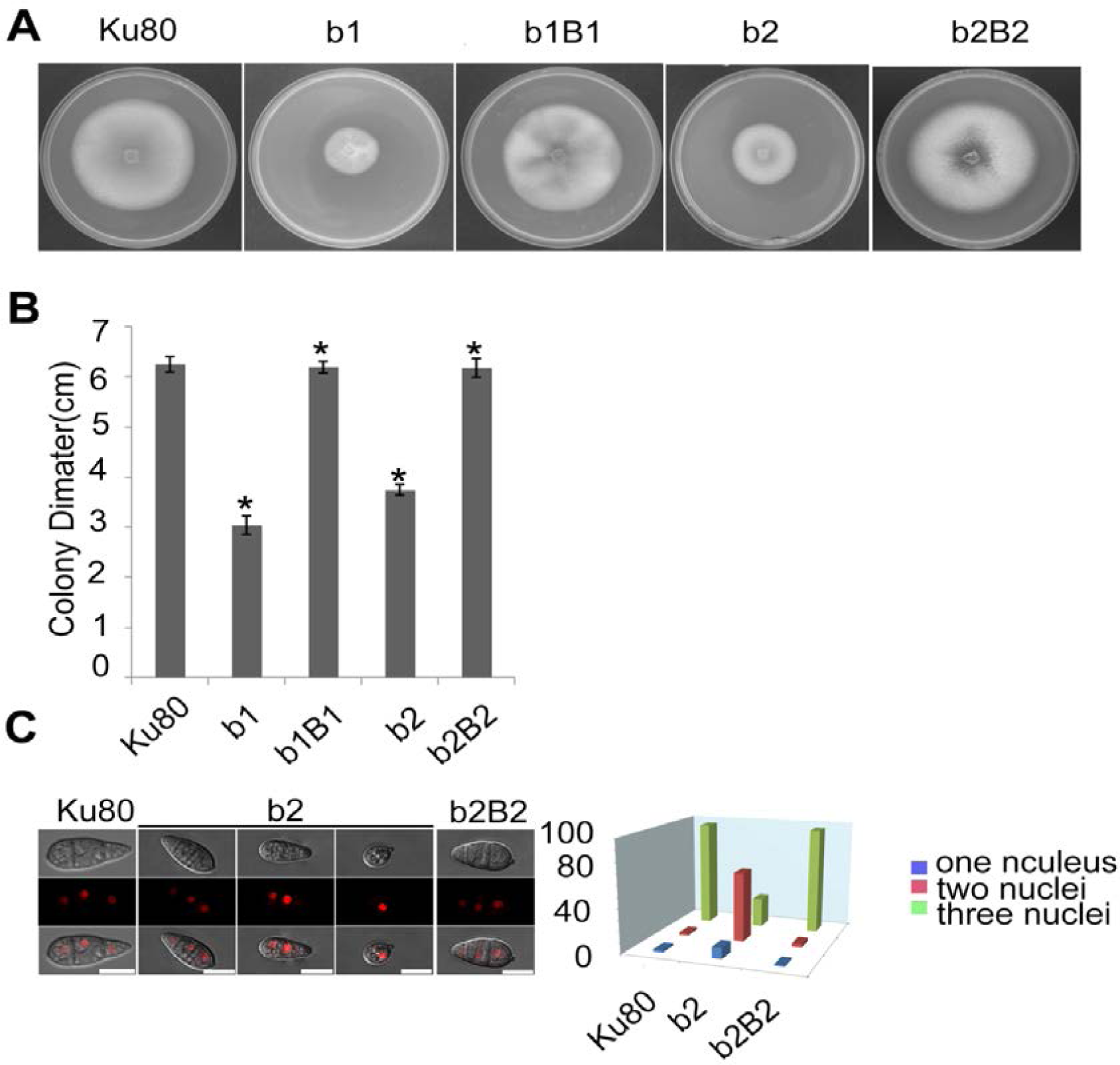
Intact CK2 holoenzyme is needed for normal growth. **A**, Colonial morphology and **B,** vegetative growth of *ΔMockb1* and *ΔMockb2* deletion mutants (b1 and b2) and their respective complementation strains (b1B1 and b2B2) was observed on SYM agar plates incubated in the dark for 10 days at 25°C, and then photographed. **C (left)** The conidial morphology of the *ΔMockb2* deletion (b2) was detected and compared to background (Ku80) and the complement (b2B2). The red fluorescence shows the nuclear number in the conidia. The red fluorescence was due to the nuclear protein histone linker (MGG_12797) fused with mCherry used as nuclear marker. All bars = 10 um. **C (right**) The percentage of conidia with different nuclear number in the conidia produced by Ku80, b2 and b2B2. Error bars shows SE and a star indicate a P<0.05 for control is same or larger than for the mutants.

### Subcellular localization of CK2 subunits

To assess the localization of the three CK2 subunits, we constructed N-terminal GFP fusions of all three proteins (Filhol et al., 2003) GFP-MoCKa, GFP-MoCKb1 and GFP-MoCKb2 (strains GfpA, GfpB1 and GfpB2 respectively **Table 1**). All three strains showed the same growth rate, morphology (data not shown) and pathogenicity (**Fig. S1A**) as the control strain Ku80. The CKa and CKb1&2 Gfp-fusion proteins localized to nuclei and prominently to nucleoli and, interestingly, to both sides of septal pores in hyphae and conidia (**Fig. 3A, C, E and J**). The RNA levels for the GFP fusion genes in these strains were elevated ∼10 to 15-fold over the control level **(Fig. S1B, C and D**).

**Fig. 3.**
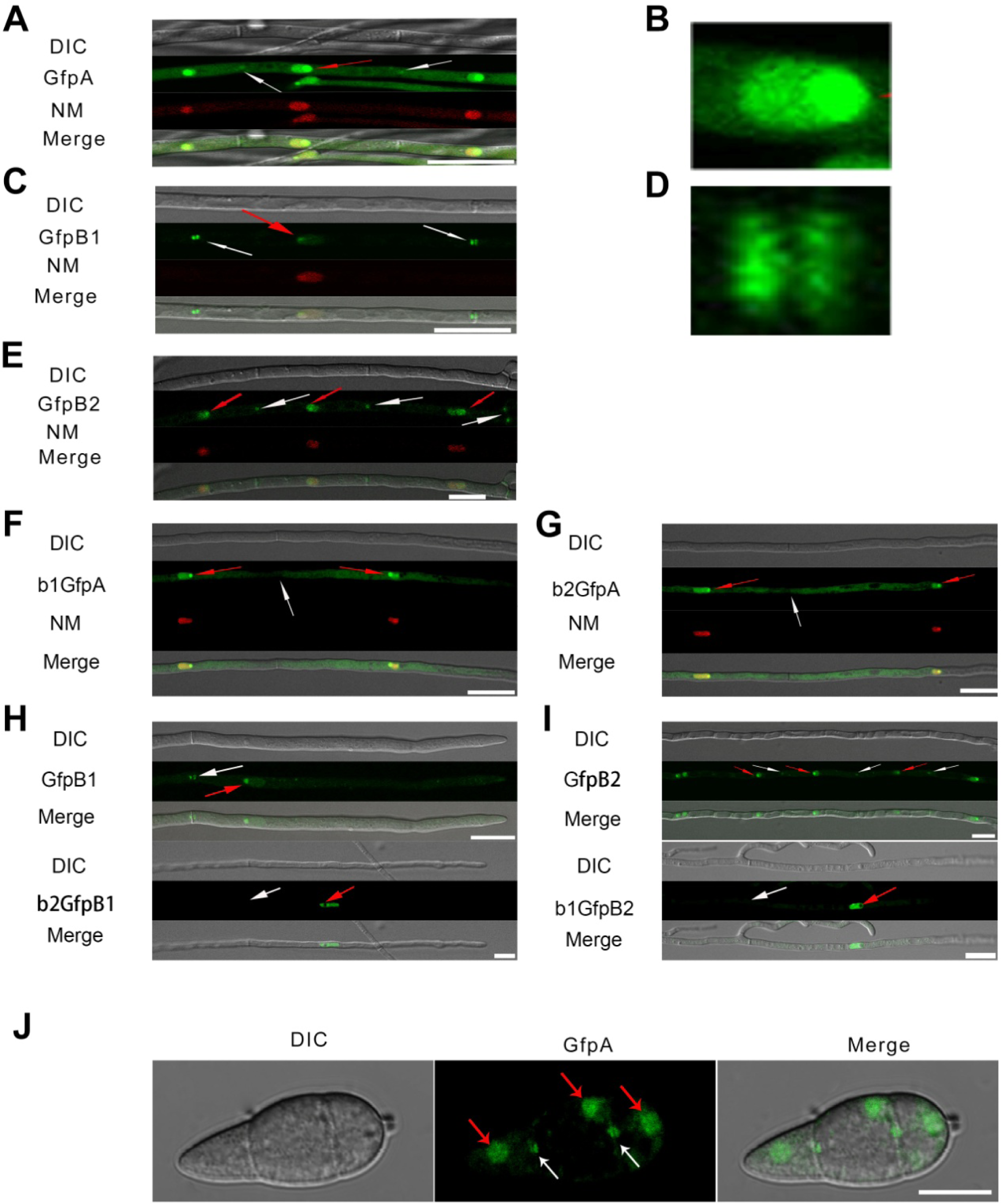
Intracellular localization of the three CK2 holoenzyme components showing that all three proteins are needed for normal localization. **A-E**, Localization of the three MoCK2 subunits in the background strain Ku80. The background strain Ku80 was transformed through gene replacements using plasmids containing GFP-MoCKa, GFP-MoCKb1 and GFP-MoCKb2 (Strains GfpA, GfpB1 and GfpB2). All three GFP constructs localize preferentially to nucleoli and to septal pores between cells. **B**, Enlargement of the nuclear localization of GFP-MoCKa (marked with red arrow in **A**). **D**, Enlargement of the septal localization of GFP-MoCKb1 (left septa marked with white arrow in **C**). **F** and **G**, Localization of over expressed GFP-MoCKa in *ΔMockb1* (b1GfpA) or in *ΔMockb2* (b2GfpA) does not rescue normal localization to septal pores. **H** and **I**, (below) Neither overexpression of MoCKb1 in *ΔMockb2* (b2GfpB1) nor overexpression of MoCKb2 in *ΔMockb1* (b1GfpB2) rescued normal localization (GfpB1 and GfpB2) (above) to nucleoli or septal pores. Histone linker (MGG_12797) was fused with the mCherry and used as nuclear localization marker (NM). **J**, GFP-MoCKa localization in conidia of strain GfpA. GFP-MoCKa localizes to nuclei (red arrows) and to septal pores (white arrows). All bars=10μm.

We then tested if the localization to septa and nucleoli were dependent on the association with the other subunits of the holoenzyme. We had measured *MoCka* expression in the two *Mockb* mutants (b1 and b2) using qPCR and noted it was downregulated two- to three-fold compared to the background control strain Ku80 (**Fig. S2A**. We constructed strains that over-express GFP-CKa in deletion strains b1 and b2 (strains b1GfpA and b2GfpA). Expression of GFP-CKa was elevated 25-fold and 15-fold in the b1GfpA and b2GfpA strains, respectively (**Fig. S2B**).

Localization to GFP-MoCKa to septa was not observed (**Fig.3F, G**), however, nucleolar localization of GFP-MoCKa was clear in the b1GfpA and b2GfpA strains. To test if over-expression of any one of the CKb proteins could rescue the effect of the deletion of the other CKb, we constructed GFP-CKb overexpression strains in both CKb mutants (strains b1GfpB2 and b2GfpB1 (**Table 1**). As noted above, the overexpression of either of the two CKbs in the control strain Ku80 showed normal localization to septa and nucleolus (**Fig.3C, E, H and I**) but the overexpression in the CKb deletion strains could not rescue normal localization (**Fig.3H and I**). Furthermore, both GFP-MoCKb1 and GFP-MoCKb2 appeared to localize to nuclei but were excluded from nucleoli in the b2GfpB1 and b1GfpB2 strains, respectively. A limited restoration of conidial production and morphology defect of *ΔMockb1&2* deletions (strains b1 and b2) was observed in b1GfpA and b2GfpA (**Fig. S2C, D and E**). In addition, a significant restoration of growth rate was detected (**Fig. 4A**).

**Fig. 4.**
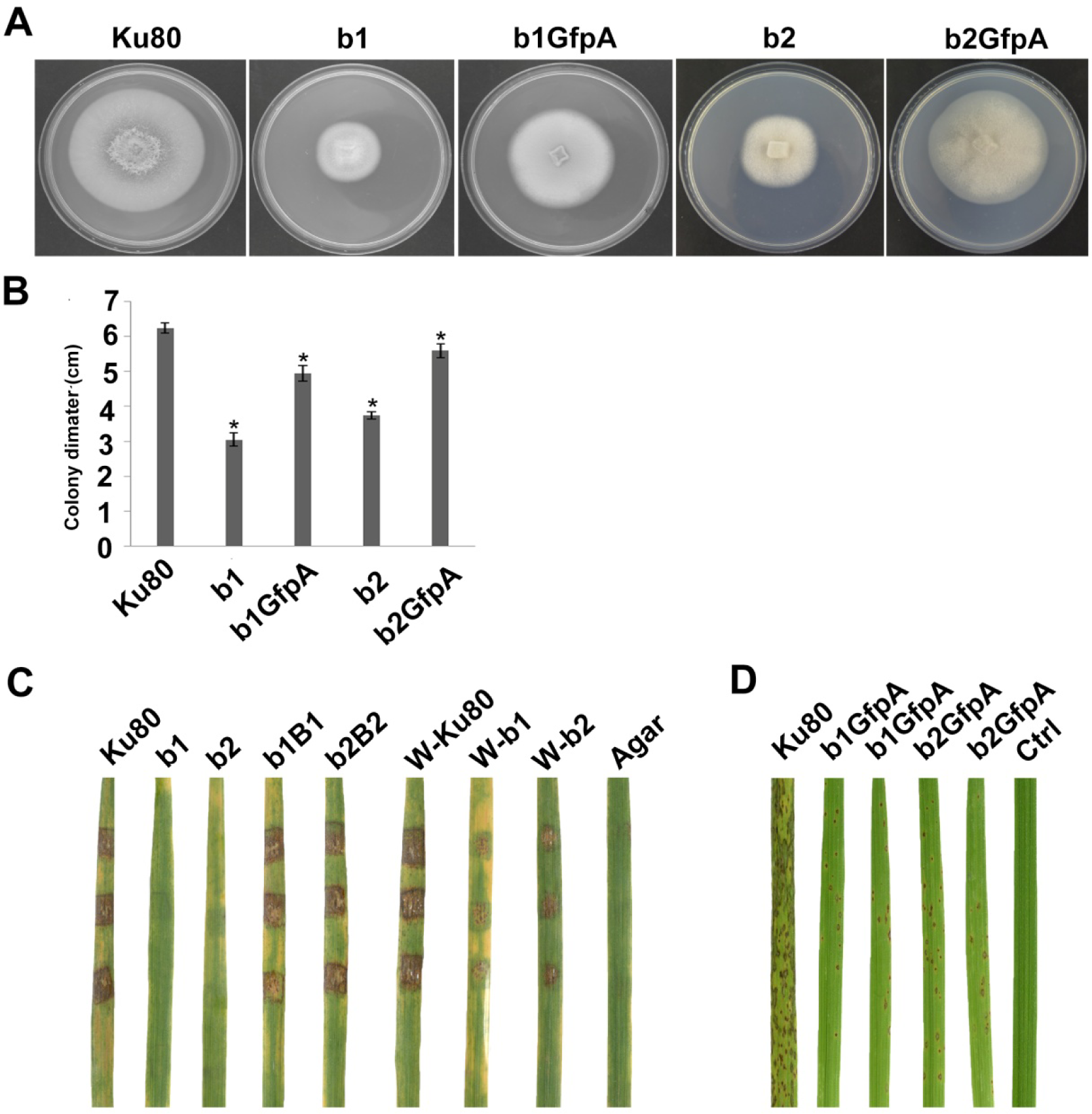
Intact CK2 holoenzyme is needed for infection and pathogenicity. (**A**) Colonial morphology and (**B**) vegetative growth of ΔMockb1, ΔMockb2, (Strain b1 and b2) and overexpressed GFP-CKa in b1 (b1GfpA) and in b2 (b2GfpA). Strains were grown on SYM medium agar plates incubated in the dark for 10 days at 25°C, and then photographed. (**C**) Pathogenicity analysis strains b1 and b2 on rice. Disease symptoms on rice leaves of 2-week-old seedlings inoculated using mycelial plugs since mutants produced no or very few conidia. Typical leaves were photographed 6 days after inoculation. The b1B1and b2B2 were complementary strains. Treatments starting with a W indicate that the leaf surface was wounded before mycelial plug was applied. The rice cultivar was CO-39. (**D**) Pathogenicity analysis of overexpressed GFP-CKa in strain b1 (b1GfpA) and b2 (b2GfpA). Disease symptoms on rice leaves of 2-week-old seedlings inoculated with conidia suspension since CKa overexpressed strains produced just enough conidia to use for the assay. The concentration of conidia in the suspension was about 1×105/ml. Typical leaves were photographed 6 days after inoculation. The rice cultivar was CO-39. Error bars shows SE and a star indicate a P<0.05 for that the control is same or larger than for the mutants.

### Infection phenotypes of CKb deletions

Deletion of CK2 genes has been shown to have effects on both growth and infection in *F. graminearum* (Wang et al., 2011) and we also found this to be the case for *M. oryzae*. Conidiation was virtually absent in both*ΔMockb1* and *ΔMockb2* deletion mutants (strains b1 and b2), thus, we used mycelial plugs to test for infection (Liu et al., 2010; Talbot et al., 1996). Compared to the background strain Ku80, mutants lacking one of the MoCK2b components had severely reduced or complete lost pathogenicity on intact leaves. However, wound inoculated leaves were impacted by the mutants (**Fig. 4C)**.

Over-expression of MoCka in the *ΔMockb1* and *ΔMockb2* lines (strains b1GfpA and b2GfpA) allowed sufficient conidia production to perform conidial inoculations. Small lesions were observed in both cases, indicating that Ckb subunits are not required for pathogenesis (**Fig. 4C**).

Overexpression of the MoCkb1 subunit in strain b2, strain b2GfpB1, restored growth rate and improved conidiation. In contrast, overexpression of MoCkb2 in strain b1 (strain b1GfpB2) did not restore growth or conidiation (**Fig. 5A**).

**Fig. 5.**
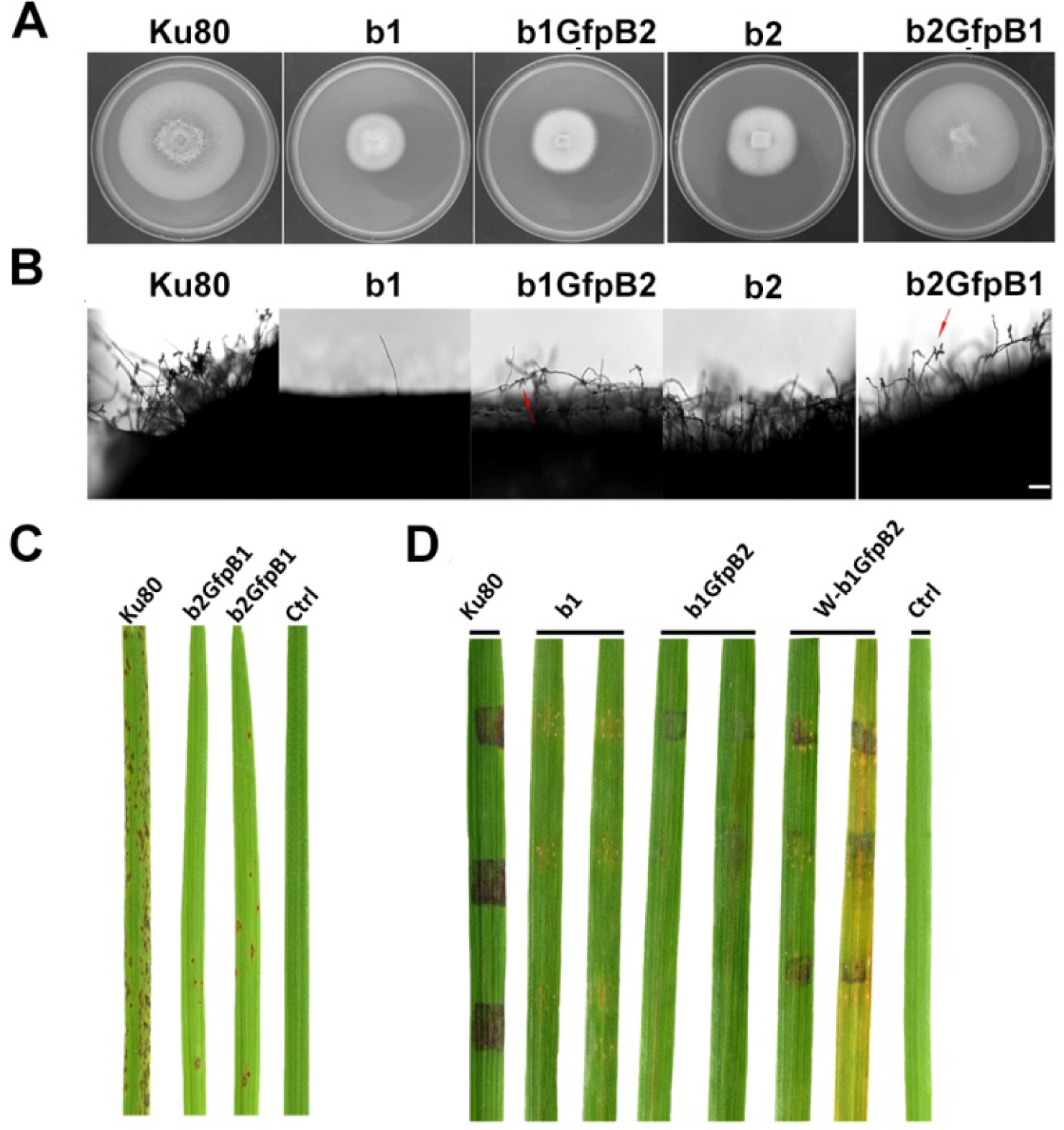
Phenotypic effects in the respective MoCKb deletion mutants of overexpression the other MoCKb component. (**A**) Colonial morphology and vegetative growth of strain b1, b1GfpB2, b2 and b2GfpB1 transformants was observed on SYM medium agar plates grown in the dark for 10 days at 25 °C and then photographed. (**B**) Development of conidia on conidiophores. Light microscopic observation was performed on strains grown on the rice bran medium for 10 days. The red arrows indicate some conidia are produced by the b1GfpB2 and b2GfpB1 strains. Bar=50um. (**C**) Pathogenic analysis of the b2GfpB1 strain. Disease symptoms on rice leaves of 2-week-old seedlings were also inoculated by conidia suspension. The concentration of conidia suspension for inoculation was about 1×10^5^/ml. (**D**) Pathogenic analysis of b1 and b1GfpB2 strains on rice. Disease symptoms on rice leaves of 3-week-old seedlings were inoculated using mycelial plugs. The W-b1GfpB2 indicates that the rice leaves were wounded before the agar plugs were placed. Typical leaves were photographed 6 days after inoculation. The rice strain was CO-39.

There was a limited but detectable restoration of pathogenicity in spore inoculation of strainb2GfpB1 compared to strain b2 and in mycelial inoculation of unwounded plants of strain b1GfpB2 compared to strain b1 (**Fig. 5C, D**). This suggests that overexpression of MoCkb2 in the *ΔMockb1* mutant was unable to compensate for vegetative functions but could partially suppress the pathogenicity defect.

### CKa localization in appressoria

Since we found large effects in infection of the deletion of the CKb components, we decided to investigate localization of GFP-CKa in the appressoria. As in hyphae and conidia, GFP-CKa (strain GfpA) localizes to nuclei (**Fig. 6A** top row), to the septa between the appressorium and the germ tube (**Fig. 6E**) and also assembles a large ring structure perpendicular to the penetration pore (**Fig. 6B, C, D,** and **Video S1-5** showing 3D rotations to visualize the ring and the appressorium). MoCKa nuclear localization was present in appressoria formed by the two CK2b deletion mutants (strains b1GfpA and b2GfpA) (**Fig. 6A** middle and bottom row), however ring structures were not observed. Concentration of GFP-MoCKa in nucleoli is clear in conidia, however, we could not clearly observe preferential nucleolar localization in appressoria. As can be seen in (**Fig. 6D**) the CK2 large ring structure is positioned perpendicular to the penetration pore where the F-actin-septin ring has been shown to form around the pore opening (Dagdas et al., 2012) (**Fig. 6D and Fig. 7**). Using the 3d image stack we measured the size of the rings seen in Fig. 3C. The left and right rings had outer diameter of 5.3 and 5.5 μm, thickness from the sides opposite and away from the penetration pore of 0.7 and 0.7 μm and thickness at the penetration pore of 1.4 and 1.2 μm, respectively.

**Fig. 6.**
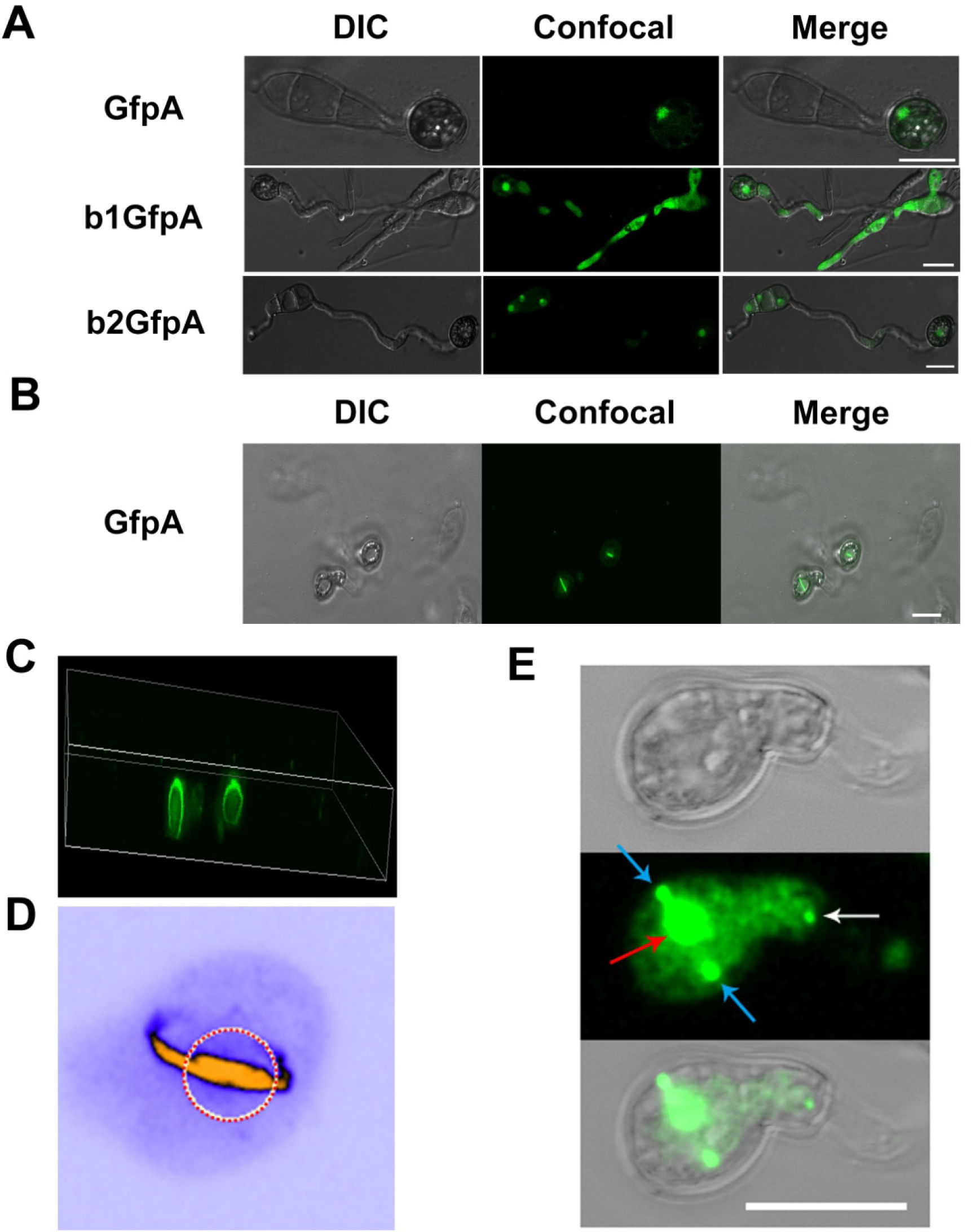
Localization of the GFP-MoCKa subunit in appressoria of the background strain Ku80 (strain GfpA) the and in the two MoCKb deletion strains (b1GfpA and b2GfpA) (compare Fig. 5A, F, G). (**A**) Localization of GFP-MoCKa in all three strains show localization to nuclei. (**B**) In the strain GfpA that form appressoria from conidia a bright line of GFP-MoCKa can be seen across the appressorium penetration pores. (**C**) Through 3d scanning and then rotating the 3d reconstruction image (**Video S 1**) we found that the streak across the penetration pores is a ring of GFP-MoCKa perpendicular to the penetration pore opening not present in the deletion strains b1GfpA and b2GfpA (**Video S2** and S**3**). (**D**) False colour lookup table 3d reconstruction image of the right ring structure in C enlarged and rotated back and seen from the same angle as in B with the penetration pore opening indicated by a red-white circle seen from the “plant” side (**Video S4** and S**5** for left and right ring in false colours). The false colour was used so that the week fluorescence in the cytoplasm could be used to show the whole cytoplasm volume. The image was made using the analytical image analysis freeware ImageJ (https://imagej.nih.gov/ij/index.html) and the ICA lookup table in ImageJ was used for false colouring. (**E**) Image in the 3d stack that shows the septal pore accumulation. Image is deliberately “overexposed” for the ring structure to be able to show nuclear and septal localization. GFP-MoCKa localizes to nuclei (red arrow) surrounded by the large ring structure in cross-section (blue arrows) and to the septal pore towards the germ tube (white arrow). All bars=10 μm

**Fig. 7.**
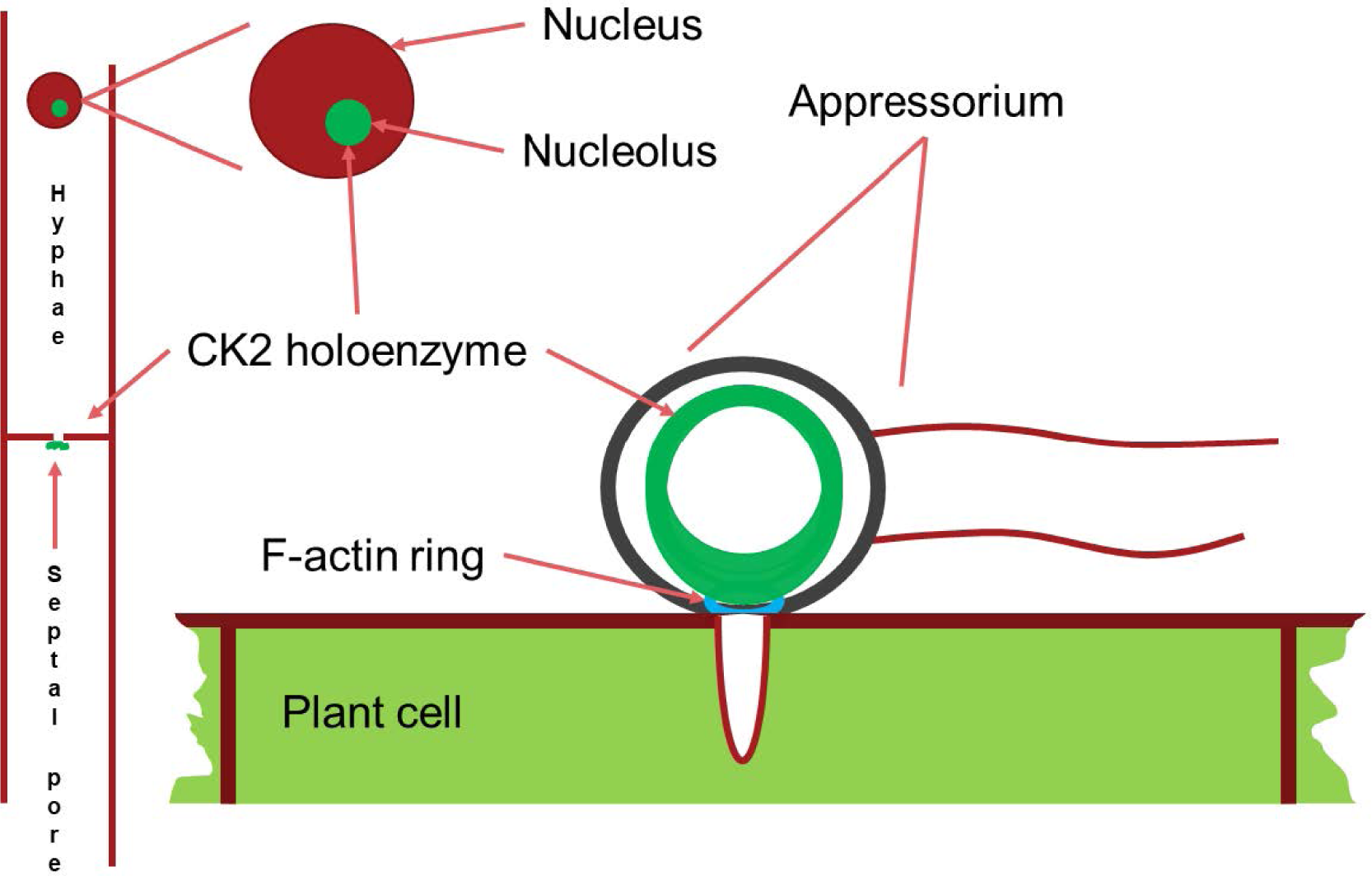
Schematic drawing of the main localizations of the CK2 holoenzyme. It localizes to the nucleolus, to septal pores between cells and forms a large ring structure perpendicular to the F-actin ring surrounding the appressorium penetration pore. Appressorium and hyphae drawn approximately to their relative sizes.

## DISCUSSION

The analysis of the *MoCKb* mutants and the localization of the GFP-labelled MoCK2 proteins showed that all identified MoCK2 components are needed for normal function and also normal localization. Localization to septa requires all three subunits, presumably as the holoenzyme. Mutation of either CKb subunit blocks nucleolar localization of the other CKb subunit. Surprisingly, nucleolar localization of CKa was observed in the CKb mutants. This shows that the holoenzyme is not required for CKa localization to the nucleolus. It seems likely that CKb1 and CKb2 must interact with each other in order to interact with CKa, and that CKa is required for movement of CKb subunits into the nucleolus as the holoenzyme.

The pattern of localization to septa (**Fig. 3**) observed is remarkably similar to that displayed by the Woronin body tethering protein AoLah from *A. oryzae* (Fig. 4b in (Han et al., 2014)). Our results thus demonstrate that the MoCK2-holoenzyme assembles as a large complex near, and is perhaps tethered to, septa, possibly through binding to MoLah. Since septal pores in fungi are gated (Shen K-F. et al., 2014), as are gap junctions and plasmodesmata in animal and plant tissue, respectively (Ariazi et al., 2017; Kragler, 2013; Neijssen et al., 2005), CK2 has a potential to play a general role in this gating.

The crystal structure of CK2 suggests that it can form filaments and higher-order interactions between CK2 holoenzyme tetramer units, and based on this it has been predicted that autophosphorylation between the units could occur to down-regulate its activity (Litchfield, 2003; Poole et al., 2005). Filament formation has been shown to occur *in vitro*(Glover, 1986; Seetoh et al., 2016; Valero et al., 1995) and *in vivo* (Hübner et al., 2014). Several forms of higher order interactions have been predicted, and it has been demonstrated that at least one of these has reduced kinase activity (Poole et al., 2005; Valero et al., 1995). However, in our localization experiments focused on septa, we cannot distinguish if the large structure is due to co-localization of the CK2 with another protein, such as the MoLah ortholog, or if CK2 is in an aggregated form near septa. Since MoLah has the characteristics of an intrinsically disordered protein (Han et al., 2014), and CK2 has been proposed to interact with proteins to promote their disordered state (Zetina, 2001; Tantos et al., 2013; Na et al., 2018), we favour the view that CK2 interacts with MoLah and other proteins to form a complex near septa. The large ring observed in appressoria (Fig. 3 & 4) may on the other hand be a true filament of CK2 in a relatively inactive state that is a store for CK2 so that upon plant penetration by the infection peg hyphae it can be activated and influence appressorial pore function, and other pathogenesis-specific functions.

Previous studies of subcytosolic localization reveals that CK2 is also associated with import into organelles. CK2 promotes protein import into endoplasmic reticulum (Wang and Johnsson, 2005) and into mitochondria during mitochondrial biogenesis and maintenance (Rao et al., 2011). CK2 phosphorylation has been shown to activate Tom22 precursors to assemble a functional mitochondrial import machinery (Rao et al., 2011).

### Conclusion

We conclude that CK2 is necessary for normal growth and plant infection by *M. oryzae.* CK2 localizes to the nucleus, in particular to the nucleolus but also to hyphal septal pores and most interestingly to appressorial penetration pores. The pore localizations are dependent on all CK2 commponents of the CK2 holoenzyme while the nuclear localization of CKa is not necessarily dependent on the presence of a specific CKb unit. These localizations of CK2 points to a possible involvement in regulating the traffic through pores and in line with this also the distribution of protein inside the nucleus.

## Supporting information

Supplemental Figures and table including captions for videos and 5 video files

## Data availability

Additional data that support the findings of this study are available from the corresponding authors upon request.

## Acknowledgements

We thank Dr. Guanghui Wang, Dr. Wenhui Zheng, Dr. Ya Li and Dr. Huawei Zheng (Fujian Agriculture and Forestry University, Fuzhou, China) for their helpful discussions. We thank Professor Jin-Rong Xu, Department of Botany and Plant Pathology Purdue University, U.S.A. for providing the Ku80 strain. We thank Dr. BjoernOest Hansen, Göttingen for help with mapping and constructing the used *M. oryzae* transcriptome data file from downloaded data. This work was supported by the National Natural Science Foundation of China (U1305211), National Key Research and Development Program of China (2016YFD0300700) and National Natural Science Foundation for Young Scientists of China (Grant No.31500118 and No.31301630), the 100 Talent Program of Fujian Province, and USDA NIFA Hatch project 1013944.

## Authors’ contributions

Conceived and designed the experiments: G. L., S. O. and Z. W. Performed the experiments: L. Z., D. Z., Y. C., W. Y. and Q. L. Analysed the data: L. Z., D. Z., D. J. E., S. O. and Z. W. Wrote the paper: L. Z., D. Z., D. J. E., S. O. and Z. W.

## Correspondence

and request for materials should be addressed to S.O. or Z.W.

## REFERENCES

Ahmad, K. A., Wang, G., Unger, G., Slaton, J., and Ahmed, K. (2008). Protein kinase CK2 – A key suppressor of apoptosis. Adv. Enzyme Regul. 48, 179–187. doi:10.1016/j.advenzreg.2008.04.002.

Ariazi, J., Benowitz, A., De Biasi, V., Den Boer, M. L., Cherqui, S., Cui, H., et al. (2017). Tunneling Nanotubes and Gap Junctions–Their Role in Long-Range Intercellular Communication during Development, Health, and Disease Conditions. Front. Mol. Neurosci. 10. doi:10.3389/fnmol.2017.00333.

Beckerman, J. L., and Ebbole, D. J. (1996). MPG1, a Gene Encoding a Fungal Hydrophobin of Magnaporthe grisea, Is Involved in Surface Recognition. MPMI 9, 450–456.

Bourett, T. M., Sweigard, J. A., Czymmek, K. J., Carroll, A., and Howard, R. J. (2002). Reef coral fluorescent proteins for visualizing fungal pathogens. Fungal Genet. Biol. 37, 211–220. doi:10.1016/S1087-1845(02)00524-8.

Dagdas, Y. F., Yoshino, K., Dagdas, G., Ryder, L. S., Bielska, E., Steinberg, G., et al. (2012). Septin-mediated plant cell invasion by the rice blast fungus, Magnaporthe oryzae. Science 336, 1590–1594.

Dean, R., Van Kan, J. A. L., Pretorius, Z. A., Hammond-Kosack, K. E., Di Pietro, A., Spanu, P. D., et al. (2012). The Top 10 fungal pathogens in molecular plant pathology: Top 10 fungal pathogens. Mol. Plant Pathol. 13, 414–430. doi:10.1111/j.1364-3703.2011.00783.x.

Ding, S.-L., Liu, W., Iliuk, A., Ribot, C., Vallet, J., Tao, A., et al. (2010). The Tig1 Histone Deacetylase Complex Regulates Infectious Growth in the Rice Blast Fungus Magnaporthe oryzae. PLANT CELL ONLINE 22, 2495–2508. doi:10.1105/tpc.110.074302.

Filhol, O., Nueda, A., Martel, V., Gerber-Scokaert, D., Benitez, M. J., Souchier, C., et al. (2003). Live-Cell Fluorescence Imaging Reveals the Dynamics of Protein Kinase CK2 Individual Subunits. Mol. Cell. Biol. 23, 975–987. doi:10.1128/MCB.23.3.975-987.2003.

Glover, C. V. (1986). A Filamentous Form of Drosophila Casein Kinase II. J. Biol. Chem. 261, 14349–14354.

Götz, C., and Montenarh, M. (2016). Protein kinase CK2 in development and differentiation (Review). Biomed. Rep. doi:10.3892/br.2016.829.

Han, P., Jin, F. J., Maruyama, J., and Kitamoto, K. (2014). A Large Nonconserved Region of the Tethering Protein Leashin Is Involved in Regulating the Position, Movement, and Function of Woronin Bodies in Aspergillus oryzae. Eukaryot. Cell 13, 866–877. doi:10.1128/EC.00060-14.

Hermosilla, G. H., Tapia, J. C., and Allende, J. E. (2005). Minimal CK2 activity required for yeast growth. Mol. Cell. Biochem. 274, 39–46. doi:10.1007/s11010-005-3112-2.

Hübner, G. M., Larsen, J. N., Guerra, B., Niefind, K., Vrecl, M., and Issinger, O.-G. (2014). Evidence for aggregation of protein kinase CK2 in the cell: a novel strategy for studying CK2 holoenzyme interaction by BRET2. Mol. Cell. Biochem. 397, 285–293. doi:10.1007/s11010-014-2196-y.

Kragler, F. (2013). Plasmodesmata: intercellular tunnels facilitating transport of macromolecules in plants. Cell Tissue Res. 352, 49–58. doi:10.1007/s00441-012-1550-1.

Litchfield, D. W. (2003). Protein kinase CK2: structure, regulation and role in cellular decisions of life and death. Biochem. J. 369, 1–15.

Liu, W., Xie, S., Zhao, X., Chen, X., Zheng, W., Lu, G., et al. (2010). A homeobox gene is essential for conidiogenesis of the rice blast fungus Magnaporthe oryzae. Mol. Plant. Microbe Interact. 23, 366–375.

Livak, K. J., and Schmittgen, T. D. (2001). Analysis of Relative Gene Expression Data Using Real-Time Quantitative PCR and the 2−ΔΔCT Method. Methods 25, 402–408. doi:10.1006/meth.2001.1262.

Meggio, F., and Pinna, L. A. (2003). One thousand-and one substrates of protein kinase CK2. FASEB J. 17, 349–368.

Na, J.-H., Lee, W.-K., and Yu, Y. (2018). How Do We Study the Dynamic Structure of Unstructured Proteins: A Case Study on Nopp140 as an Example of a Large, Intrinsically Disordered Protein. Int. J. Mol. Sci. 19, 381. doi:10.3390/ijms19020381.

Neijssen, J., Herberts, C., Drijfhout, J. W., Reits, E., Janssen, L., and Neefjes, J. (2005). Cross-presentation by intercellular peptide transfer through gap junctions. Nature 434, 83–88.

Padmanabha, R., Chen-Wu, J. L.-P., Arnot, D. E., and Glover, C. V. C. (1990). Isolation, sequencing, and disruption of the yeast CKA2 gene casein kinase II is essential for viability in Saccharomyces cerevisiae. Mol. Cell. Biol. 10, 4089–4099.

Poole, A., Poore, T., Bandhakavi, S., McCann, R. O., Hanna, D. E., and Glover, C. V. (2005). A global view of CK2 function and regulation. Mol. Cell. Biochem. 274, 163–170.

Rao, S., Gerbeth, C., Harbauer, A., Mikropoulou, D., Meisinger, C., and Schmidt, O. (2011). Signaling at the gate: Phosphorylation of the mitochondrial protein import machinery. Cell Cycle 10, 2083–2090. doi:10.4161/cc.10.13.16054.

Seetoh, W.-G., Chan, D. S.-H., Matak-Vinković, D., and Abell, C. (2016). Mass Spectrometry Reveals Protein Kinase CK2 High-Order Oligomerization via the Circular and Linear Assembly. ACS Chem. Biol. 11, 1511–1517. doi:10.1021/acschembio.6b00064.

Shen K-F., Osmani A. S., Govindaraghavan M., and Osmani S. A. (2014). Mitotic rregulation of fungal cell-to-cell connectivity through septal pores involves the NIMA kinase. Mol. Biol. Cell 25, 763–775. doi:10.1091/mbc.E13-12-0718.

Shlezinger, N., Goldfinger, N., and Sharon, A. (2012). Apoptotic-like programed cell death in fungi: the benefits in filamentous species. Front. Oncol. 2. doi:10.3389/fonc.2012.00097.

Sweigard, J. A., Chumley, F., Carroll, A., Farrall, L., and Valent, B. (1997). A series of vectors for fungal transformation. Fungal Genet. Rep. 44, 52–53. doi:10.4148/1941-4765.1287.

Talbot, N. J., Kershaw, M. J., Wakley, G. E., De Vries, O. M., Wessels, J. G., and Hamer, J. E. (1996). MPG1 encodes a fungal hydrophobin involved in surface interactions during infection-related development of Magnaporthe grisea. Plant Cell Online 8, 985–999.

Tantos, A., Szrnka, K., Szabo, B., Bokor, M., Kamasa, P., Matus, P., et al. (2013). Structural disorder and local order of hNopp140. Biochim. Biophys. Acta BBA - Proteins Proteomics 1834, 342–350. doi:10.1016/j.bbapap.2012.08.005.

Valero, E., De Bonis, S., Filhol, O., Wade, R. H., Langowski, J., Chambaz, E. M., et al. (1995). Quaternary Structure of Casein Kinase 2 - Characterization of Multiple Oligomeric States and Relation with its Catalytic Activity. J. Biol. Chem. 270, 8345–8352.

Villalba, F., Collemare, J., Landraud, P., Lambou, K., Brozek, V., Cirer, B., et al. (2008). Improved gene targeting in Magnaporthe grisea by inactivation of MgKU80 required for non-homologous end joining. Fungal Genet. Biol. 45, 68–75. doi:10.1016/j.fgb.2007.06.006.

Wang, C., Zhang, S., Hou, R., Zhao, Z., Zheng, Q., Xu, Q., et al. (2011). Functional Analysis of the Kinome of the Wheat Scab Fungus Fusarium graminearum. PLoS Pathog. 7, e1002460. doi:10.1371/journal.ppat.1002460.

Wang, X., and Johnsson, N. (2005). Protein kinase CK2 phosphorylates Sec63p to stimulate the assembly of the endoplasmic reticulum protein translocation apparatus. J. Cell Sci. 118, 723–732. doi:10.1242/jcs.01671.

Zetina, C. R. (2001). A Conserved Helix-Unfolding Motif in the Naturally Unfolded Proteins. Proteins Struct. Funct. Genet. 44, 479–483.

Zhao, X., Xue, C., Kim, Y., and Xu, J.-R. (2004). A Ligation-PCR Approach for Generating Gene Replacement Constructs in Magnaporthe grisea. Fungal Genet. Rep. 51, 17–18. doi:10.4148/1941-4765.1137.

Zheng, W., Chen, J., Liu, W., Zheng, S., Zhou, J., Lu, G., et al. (2007). A Rho3 Homolog Is Essential for Appressorium Development and Pathogenicity of Magnaporthe grisea. Eukaryot. Cell 6, 2240–2250. doi:10.1128/EC.00104-07.

