## Supplemental Figures and table including captions for videos and 5 video files for "*Magnaporthe oryzae* CK2 is involved in rice blast pathogenesis and accumulates in nuclei, nucleoli, at septal and appressoria pores and forms a large ring structure in appressoria": Supplementary information and captions for Videos.pdf

**SUPPLEMENTARY TABLES AND FIGURES**

Table S1 Primers used in this study

| Primer name | The sequence of primer（5’ → 3’） |
|---|---|
| 3696qRTF | CGTCAACTACCAGAAATGCG |
| 3696qRTR | TGACGGAGTCTTGCTCTGTG |
| 446qRTF | GCAGAGGTGTCGGAGGAAT |
| 446qRTR | CCAAGATCATCTCCAGTGCC |
| 5651qRTF | ACCCGTTGCTGCCGATGG |
| 5651qRTR | TAGACCTGGAAGAGGATGTTGTGG |
| Tub1RTF | CAACATCCAGACCGCTCTC |
| Tub1RTR | ACCGACACGCTTGAACAG |
| 446AF | GCCCAACCTTTCATCCTA |
| 446AR | TTGACCTCCACTAGCTCCAGCCAAGCCTACCTCCAGTGCCTCCTT |
| 446BF | GAATAGAGTAGATGCCGACCGCGGGTTCTCGTCCAACTCTAAACTAAC |
| 446BR | GCTGGGTAAACATCTCATT |
| 5651AF | GGGGTACCCCCTCTAAGTGGTCGTGC |
| 5651AR | CCGGAATTCCTTGGATGGAATTGTGCC |
| 5651BF | CGCGGATCCAGGGAGGCGTTATCATTTA |
| 5651BR | TAATCTAGACAGAGCCGAGCTTGTCTA |
| 446comF | GCTCTAGAGCGGAACCAGTAGTTGACGG |
| 446comR | GGGGTACCCATGACAACGCCGGAGGG |
| 5651comF | GCTCTAGAGCCCGACAAGCACAAAAGAT |
| 5651comR | CCCCCGGGGAGCGTTCGTTTAGACCC |

| | |
|---|---|
| 3696GFPF | CGGGATCCATGCACAGCATGGCACGC |
| 3696GFPR | CGGAATTCTGTTGAAATTACCAGCGATTC |
| 5651GFPF | CGGGATCCATGGAAGATTTTGGCAGCG |
| 5651GFPR | CCCTCGAGTCAGACACCTTGCATCATG |

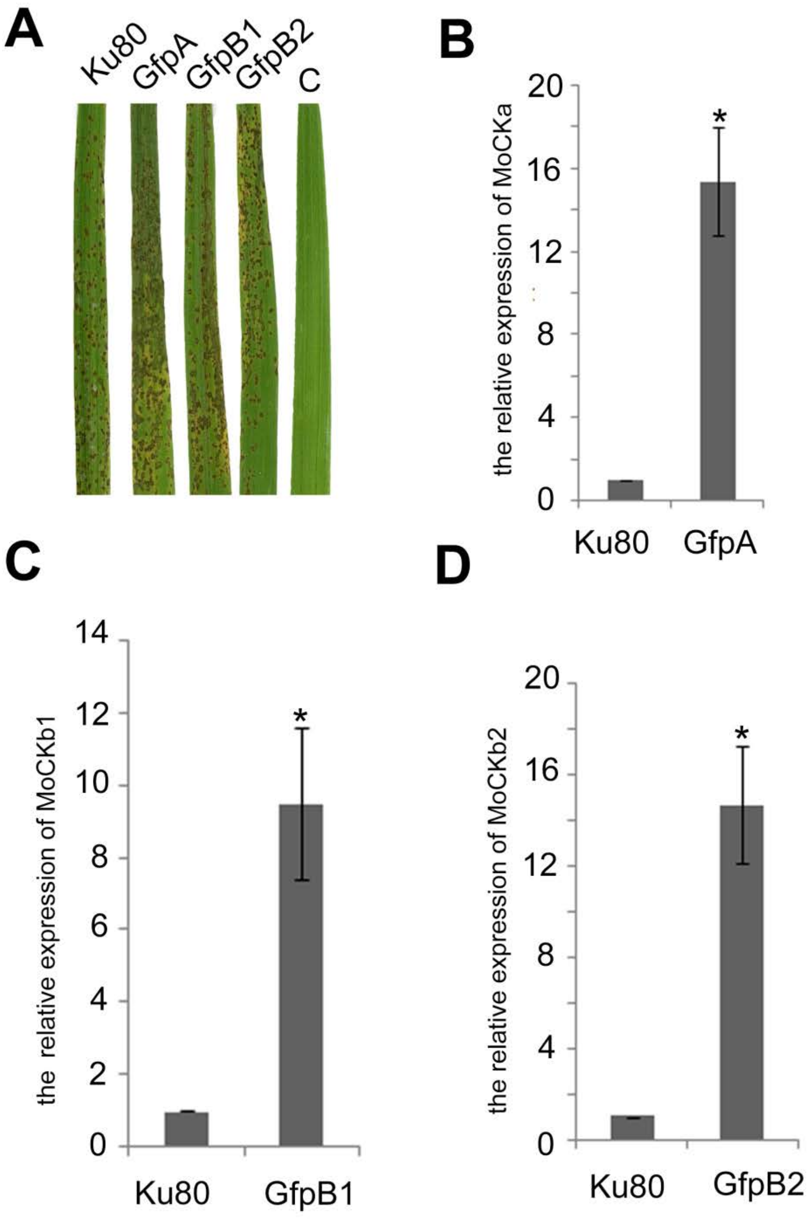


**Figure S1**. (A) Control that strains GfpA, GfpB1 and GfpB2 are as pathogenic as the background Ku80. C is untreated control. (B, C and D) the relative expression of MoCKa, MoCKb1 and MoCKb2 in the GfpA, GfpB1 and GfpB2 strains, respectively.

**A**

| Expression of | Control | Test | Test/control | SE (N=9) | P=same or higher than control |
|---|---|---|---|---|---|
| MoCKa | Ku80 | b1 | 0.38 | 0.01 | 2.07E-11 |
| MoCKa | Ku80 | b2 | 0.54 | 0.05 | 5.12E-06 |

**B**

| Expression of | Control | Test | Test/control | SE (N=9) | P=same or lower than control |
|---|---|---|---|---|---|
| MoCKa | b1 | b1GfpA | 25.11 | 1.32 | 4.23E-08 |
| MoCKa | b2 | b2GfpA | 14.60 | 1.45 | 6.85E-06 |

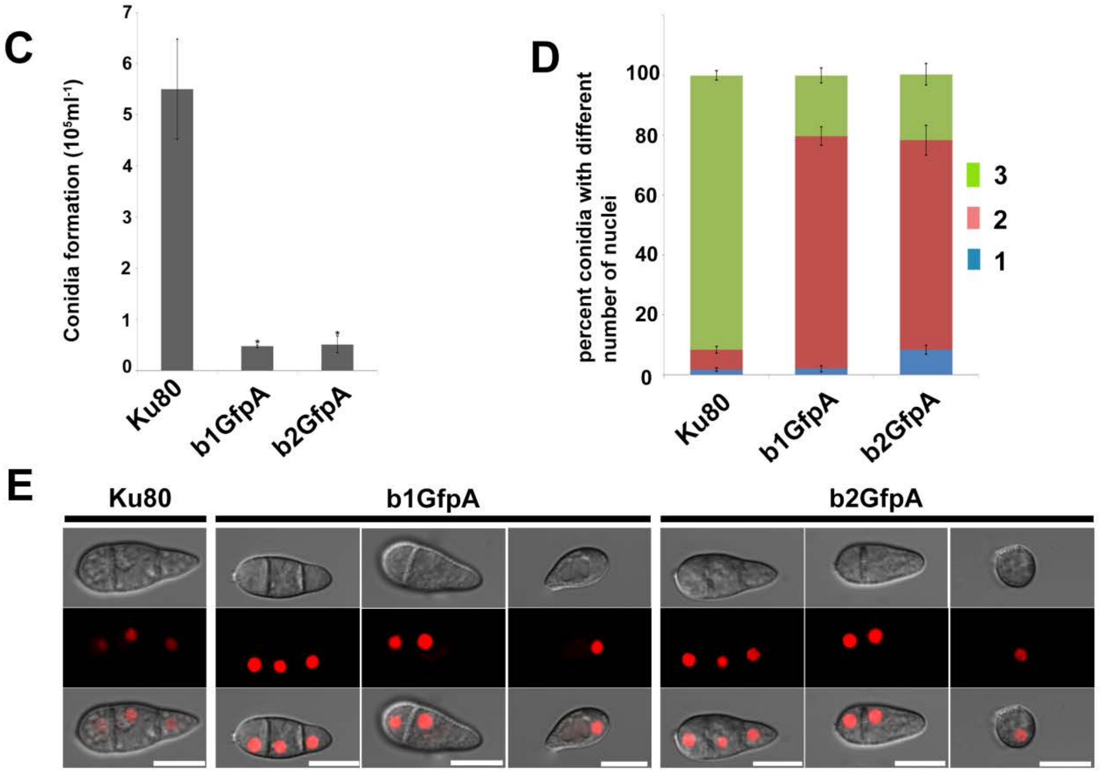


**Figure S2.** Overexpression of MoCKa in the MoCKb deletion mutants and the effect of this on conidia morphology. (**A**) MoCKa expression in the ΔMockb1 and ΔMockb2 deletion strains (b1 and b2) relative to the control Ku80 showing that expression of the other CKb were both reduced in the CKb deletion mutant. (**B**) The relative expression of MoCKa in the b1GfpA and b2GfpA in relation to their respective control backgrounds b1 and b2. (**C**) The conidial forming ability of the transformant strains b1GfpA and b2GfpA compared to the background strain Ku80. (**D**) The percentage of conidia with different numbers of nuclei produced by the background strain Ku80 and the b1GfpA and b2GfpA strains and (**E**) the conidia morphology of the three strains.

The red fluorescence was due to the nuclear protein histone linker (MGG_12797) fused with the mCherry used as nuclear marker. All bars = 10 μm.

CAPTIONS FOR SUPPLEMENTARY FILES

Video 1. By 3d scanning and then rotating the 3d reconstruction image we found that the streak across the penetration pores is a ring of GFP-MoCKa perpendicular to the penetration pore opening

Video 2. A ring as seen in Video 1 is not present in appressoria of the deletion strain b1GfpA

Video 3. A ring as seen in Video 1 is not present in appressoria of the deletion strain b2GfpA

Video 4 False color lookup table 3d reconstruction image of the left ring structure in Figure3C and Video 1.

Video 5. False colour lookup table 3d reconstruction image of the right ring structure in Figure3C and Video 1
